# An Integrated Optogenetic and Bioelectronic Platform for Regulating Cardiomyocyte Function

**DOI:** 10.1101/2023.12.15.571704

**Authors:** Olurotimi A. Bolonduro, Zijing Chen, Yan-Ru Lai, Megan Cote, Akshita A. Rao, Haitao Liu, Emmanuel S. Tzanakakis, Brian P. Timko

## Abstract

We report an integrated optogenetic and bioelectronic platform for stable and long-term modulation and monitoring of cardiomyocyte function in vitro. Optogenetic inputs were achieved through expression of a photoactivatable adenylyl cyclase (bPAC), that when activated by blue light caused a dose-dependent and time-limited increase in autonomous cardiomyocyte beat rate. Bioelectronic readouts were achieved through an integrated planar multi-electrode array (MEA) that provided real-time readouts of electrophysiological activity from 32 spatially-distinct locations. Irradiation at 27 μW/mm^2^ resulted in a ca. 14% increase in beat rate within 20-25 minutes, which remained stable for at least 2 hours. The beating rate could be cycled through repeated “on” and “off” states, and its magnitude was a monotonic function of irradiation intensity. Our integrated platform opens new avenues in bioelectronic medicine, including closedloop feedback systems, with potential applications for cardiac regulation including arrhythmia diagnosis and intervention.

**Teaser:** A system that integrates optogenetic stimulation and bioelectronic recording capabilities allows for on-demand regulation of cardiac cell function.

## Introduction

Bioelectronic medicine (BM) is an emerging field aimed at treating pathologies with devices that modulate the activity of electrically-active cells. As a distinct departure from conventional pharmacological interventions, these therapies can be localized to a particular region of the body and provide stimuli that are personalized to the patient, potentially improving clinical outcomes (*1, 2*). Platforms that can both modulate and monitor tissue functions may offer further advantages through closed-loop feedback that could intervene in the event of an acute anomaly, or adapt the therapy should baseline activity change over time. BM could be especially transformative in cardiology, where devices implanted onto the vagus nerve or the heart could address indications such as cardiac arrythmia, atrial fibrillation and cardiomyopathy (*3-5*). BM techniques that address these challenges, however, will require new classes of devices that seamlessly integrate with the surrounding tissue, causing minimal inflammation at chronic time points.

Bioelectronic recording elements such as multi-electrode arrays (MEAs) and field-effect transistors are especially attractive in BM (*6, 7*). These devices couple with electrically-active cells to provide continuous, real-time readouts of spikes that correlate with action potentials. They play an important role in cardiac tissue engineering, as biopolymer scaffolds with MEAs or multiplexed transistors have been integrated with 3D tissue constructs to interrogate tissue development and electrical synchrony (*8-11*). These devices cause minimal disruption to the cell membrane and do not elicit pronounced immune response (*12*), making them uniquely suitable for long-term tissue integration. Bioelectronic scaffolds within cardiac organoids have tracked organogenesis for 35 days (*13*), while similar devices implanted in a rodent model provided readouts from the same set of cells for over one year (*14*). Their inherent stability makes bioelectronics especially useful for monitoring cardiac tissue function and identifying abnormal states. For example, we recently demonstrated a heart-on-a-chip system that could monitor electrophysiological responses to acute hypoxia (*15*). Soft and bioresorbable MEAs implanted onto the heart in a rodent model may provide diagnostic readouts of organ-level electrophysiology (*16*).

Optogenetic techniques represent one promising path toward modulating cellular function with high spatial and temporal resolution and without generating electrical artifacts that convolute bioelectronic recording. Microbial opsins have emerged as optically-transduced modulators of transmembrane trafficking of ions and have been widely explored in neuroscience (*17, 18*). Opsins chimerized with G protein-coupled receptors (GPCR, OptoXRs) were shown to change the intracellular concentration of secondary messengers thereby broadening the repertoire of optogenetic applications to non-excitable cells. The potential of optogenetics has prompted advances in light-mediated modulation of cellular function including vectors which target specific cell types or subcellular compartments, multi-modal light sources for independent control over different optogenetic moieties, and fiber optic arrays to control tissue function *in vivo* (*17, 18*). Within the past decade these techniques have been extended to cardiac systems, with potential applications for arrythmia management and pacing, as a safer alternative to current pacemaker technologies (*19*).

The activity of adenylyl cyclases (ACs), which convert adenosine triphosphate (ATP) into cAMP, has been a target for optogenetic modulation (*20*). Methods to regulate intracellular cAMP ([cAMP]_i_) using light would be relevant to a wide variety of cell types given the ubiquity of this second messenger. In the case of cardiomyocytes (CMs), cAMP phosphorylates protein kinase A (PKA) which in turn drives excitation-contraction coupling (*21*) involving L-type Ca^2+^ channels, phospholamban, ryanodine receptors, and troponin I (*22*). In addition, cAMP stimulates hyperpolarization-activated cyclic nucleotide-gated (HCN) channels influencing the basal beating rate of CMs (*23, 24*). As expected, rapid degradation of cAMP by phosphodiesterases (*25*) also alters [cAMP]_i_ providing time-limited control over cardiac contractility.

Optical modulation of [cAMP]_i_ in CMs may be achieved using photoactivatable ACs (PACs), which are native to microorganisms such as *Euglena gracilis* (*26*), *Oscillatoria acuminata* (*27*), and *Beggiatoa* (bPAC) (*28, 29*), or engineered (*30, 31*). The small protein (350 aa) bPAC is a homodimer of an N-terminal blue light using flavin (BLUF) photoreceptor domain and a C-terminal type III AC domain (*28*). The flavin adenine dinucleotide needed for BLUF activity is readily available in animal cells. Activation of the AC site results from conformational changes in the BLUF domain brought about by exposure to blue light. Compared to other PACs, bPAC displays lower dark activity, up to 300-fold light/dark activity, superior solubility, and short half-life of the active lit state (*30, 32, 33*). We recently expressed bPAC in murine pancreatic β-islet cells and found that photostimulation rapidly elevated [cAMP]_i_ with concomitant increase in insulin secretion (*34*). Transplantation of bPAC-expressing β-cells into diabetic mice led to improved glucose tolerance and lowered hyperglycemia upon illumination (*35*).

Herein we report an engineered cardiac system with two-way communication capabilities enabled by (a) time- and dose-modulated blue light as an input, and (b) multiplexed, real-time bioelectronic recordings as outputs (Fig. 1). The bPAC was expressed for the first time in CMs to test our hypothesis that optical inputs would elevate [cAMP]_i_ and increase the beat rate (*36*). We monitored CM activity via an integrated 32-element MEA that provided continuous readouts in both light and dark states. The outputs allowed us to monitor the dynamic changes in beating rate in response to time-limited and dose-modulated periods of irradiation. Given the ubiquity of both MEAs and light sources (e.g., light-emitting diodes) in bioelectronics, our integrated platform opens new avenues for closed-loop control of function in cardiac or other tissues.

**Fig. 1.**
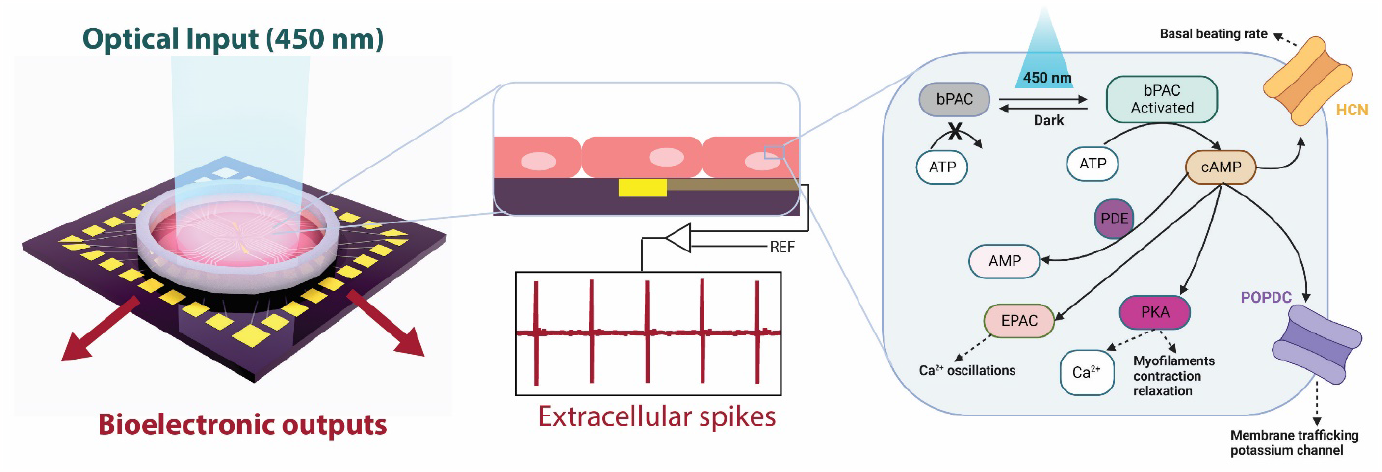
Overview of optical / bioelectronic control concept. (left) Bioelectronic chip featuring optical input and multiplexed bioelectronic outputs. (center) Detail of (red) CMs interfaced with (yellow) a single bioelectronic recording element. (right) Proposed signaling pathway in bPAC-transduced CMs whereby optical irradiation stimulates cAMP synthesis and increases beating rate. (d) Representative bioelectronic outputs.

## Results

### AdbPAC regulates [cAMP]_i_ expression with illumination

We first sought to determine whether expression of bPAC in CMs was possible. Primary rat CMs were transduced with an adenovirus (*34*) carrying a cassette with the bPAC featuring a C-terminal c-Myc epitope for immunodetection and the fluorescent reporter mCherry (AdbPAC; Fig. 2A). Western blot analysis of transduced CMs revealed a band at ca. 41 kDa, consistent with the expected size of bPAC (Fig. 2B). No expression was observed in cells with no transduction (NT) or transduced with an adenovirus (AdGFP) carrying the green fluorescent protein (GFP) gene.

**Fig. 2.**
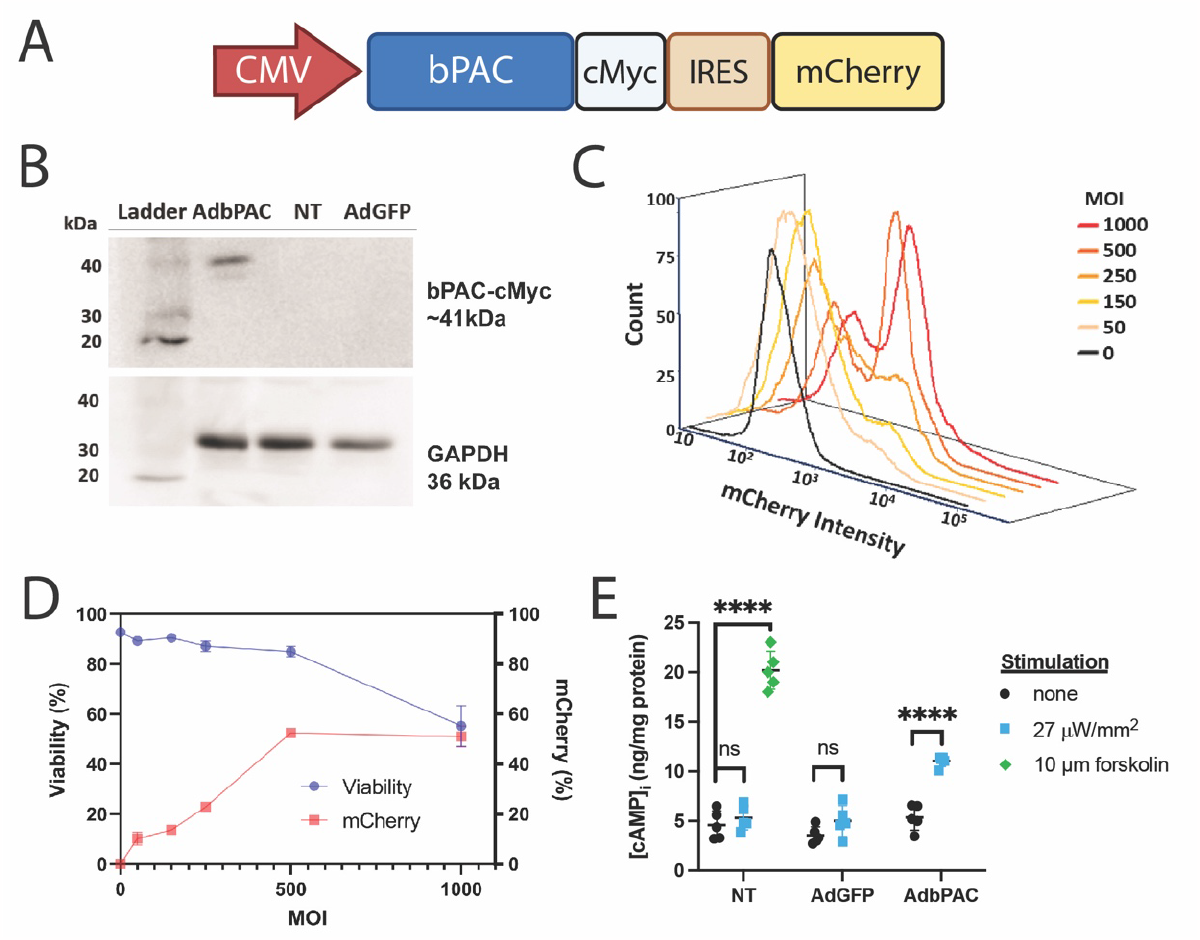
Optical interfacing using bPAC. (A) Schematic of the bPAC cassette used for adenoviral transduction of CMs. (B) Western blot analysis of samples from NT, GFP- and bPAC-expressing CMs: Top panel: bPAC (ca. 41 kDa; detection of c-Myc tag). Bottom: GAPDH (36 kDa; loading control). (C) Flow cytometry for mCherry expression in CMs with MOI ranging between 0 and 1000. The black curve corresponding to MOI of 0 (NT), represents background fluorescence. (D) Viability and mCherry expression in CMs 3 days post-transduction with the same MOIs as in (C). N=3. (E) Intracellular cAMP expression in NT, GFP^+^ and bPAC^+^ CMs without (black) or with 30-minute illumination (blue), or 10 μM forskolin (green). Comparisons represent two-tailed t-test. Non-stimulated groups have no significant difference by ANOVA. N=5 for all groups, ****P<0.0001.

After successfully expressing bPAC in CMs, flow cytometry was employed to determine the fraction of bPAC^+^ cells at various multiplicities of infection (MOI) (Fig 2C). The expression of mCherry was positively correlated with MOI over the range of 0-500 but plateaued between 500-1000 with expression at 52±1.0% and 51±4.2% at those two endpoints, respectively (Fig. 2D). Increasing the MOI from 500 to 1000 also decreased the cell viability from 85.0±2.1% to 55.3±8.2%. For these and all subsequent experiments, MOI was set to 500.

Next, the modulation was studied of [cAMP]_i_ with light stimulation in bPAC-expressing CMs (Fig. 2E). The cells were illuminated with a 450 nm light emitting diode (LED) array continuously for 30 minutes. Here and for all subsequent experiments, unless noted otherwise, an intensity of 27 μW/mm^2^ was employed. Compared to AdbPAC CMs kept in the dark, irradiated cells exhibited nearly a 2-fold increase in [cAMP]_i_ (11.04±0.49 vs. 5.36±1.15 ng cAMP/mg total protein, p<0.0001, n = 5). By contrast, illumination did not significantly change [cAMP]_i_ in NT or GFP-expressing CMs. In addition, no significant difference was observed between all three CM groups without exposure to blue light, supporting that bPAC has minimal dark activity and transduction alone did not alter [cAMP]_i_ expression. Treatment with 10 µM forskolin, an activator of AC (*37*), resulted in a 1.8 -fold greater [cAMP]_i_ in NT CMs compared to AdbPAC CMs stimulated with light (20.20±1.92 vs. 11.04±0.49 ng cAMP/mg total protein, p<0.0001, n=5). This discrepancy is consistent with observations in *Xenopus laevis* oocytes (*38*), and likely arises because forskolin enhances the activity of multiple native AC isoforms whereas light activates only bPAC.

### Bioelectronic recording elements form stable interfaces with bPAC-expressing CMs

We hypothesized that multi-electrode arrays (MEAs) would stably interface with and record signals from CMs expressing bPAC. We fabricated our MEAs on silicon/silicon oxide substrates by photolithography (*15*). Each chip consisted of 32 Au-pad recording elements, two large-area quasi-reference electrodes and associated interconnects. The recording elements had a pitch of 200 μm and the full array could interrogate a 1 mm^2^ region of cell monolayer (Fig. 3A). This area is sufficiently large to calculate wavefront propagation velocities, and to identify potential variations in intercellular signaling as we observed in hypoxia studies (*15*). Each chip was mounted onto a custom printed circuit board (PCB) and the cell culture area was defined by a silicone well adhered with poly(dimethyl siloxane) (PDMS) (Fig. S1). Recording elements were electrochemically coated with platinum black to increase signal-to-noise ratio by reducing impedance ca. 32x to 8.0 kΩ at 1 kHz (Fig. S2).

**Fig. 3.**
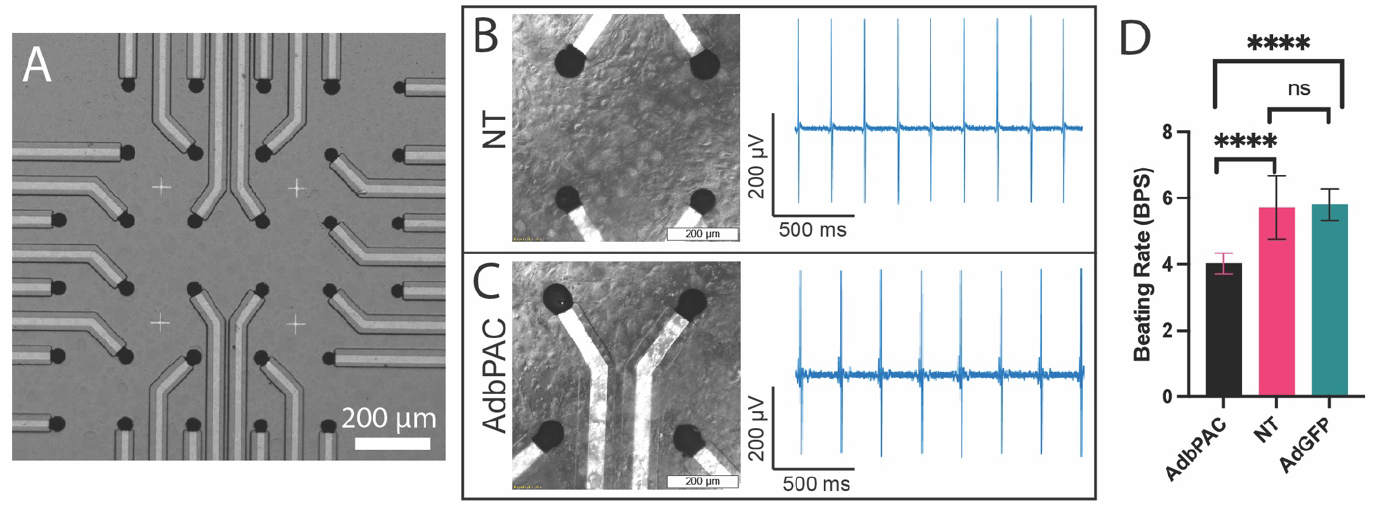
Bioelectronic interfaces with MEAs. (A) Optical image of the MEA with 32 recording elements coated with platinum black. (B,C) Representative bioelectronic interfaces with NT and bPAC-expressing CMs: (left) Representative phase contrast images and (right) corresponding bioelectronic readouts at 7 DIV. Scale bar: 30 μm. (D) Beat rate comparison between AdbPAC, NT and AdGFP groups at 6-7 DIV; N=21, N=9 and N=5, respectively. ****P<0.0001 by one-way ANOVA.

For all bioelectronics studies, CMs were plated onto MEAs coated with gelatin-fibronectin to facilitate adhesion. The CMs formed confluent monolayers that began beating as soon as 24 hours in vitro. Transduced groups were infected at 3 days in vitro (DIV) and maintained in standard culture conditions thereafter. At 6 DIV and beyond both NT- and AdbPAC-CM cultures continued to beat spontaneously and were unperturbed by the underlying devices (Fig. 3B,C). Bioelectronic readouts demonstrated periodic beating with signal-to-noise as high as 45 dB which was similar to our previous studies on HL-1 cell monolayers (*15, 39*).

At 6-7 DIV we found that NT CMs beat with a rate of 5.7 ± 1.0 beats per second (BPS) across 9 distinct cultures, consistent with the range of values reported for similar cultures of primary CMs (*40*). Cardiomyocytes transduced with AdGFP beat at a statistically similar rate, 5.8 ± 0.5 BPS (p=0.84), demonstrating that the transduction process did not significantly alter processes associated with contraction. Cells expressing bPAC, however, beat at a slower rate of 4.3 ± 0.6 BPS (compared to NT, p<0.0001) (Fig. 3D).

To further investigate the interface between cells and the chip surface we performed immunofluorescence studies on CMs cultured on glass surfaces (Fig. 4). Both NT and AdbPAC CMs exhibited features consistent with healthy cultures, including well-defined nuclei (DAPI), punctate gap junction expression at perinuclear and cell-to-cell adhesion sites based on staining of connexin-43 (Cx43), and sarcomeres with well-defined z-disks (α-actinin) characteristic of contractile CMs. In the case of CMs transduced with AdbPAC we also observed fluorescence throughout the cytosol, consistent with the expected distribution of the soluble mCherry tag.

**Fig. 4.**
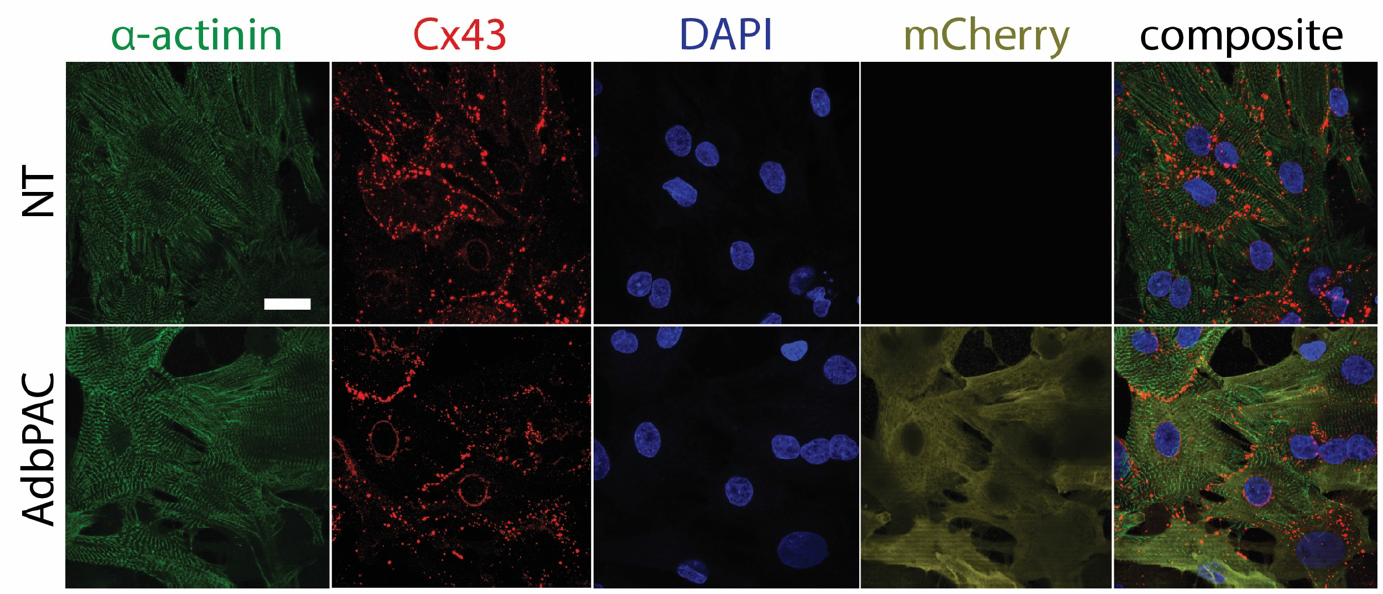
Immunofluorescence. Images of NT and bPAC-expressing (AdbPAC) CMs showing α-actinin cytoskeleton, Cx-43 gap junction proteins, nuclei, and mCherry. Scale bar: 20 μM.

### Optical and bioelectronic interfaces enable two-way communication with CMs

With the bioelectronic system established, integrating the cellular and electronic components for proper interfacing, the signal readouts were used to evaluate the effect of photostimulation on CM function (Fig. 5A,B). For the first set of assays, we measured readouts from CMs in the dark state for five minutes, followed by illumination for 20 minutes. We abstracted the beating rate (beats per second, BPS) from the electrophysiological readouts and normalized it as the fold change to the initial, basal beating rate (BPS/BPS_0_), to account for variability between cultures. We observed a progressive increase in beating rate in the bPAC group, with a 14% increase by the end of the irradiation period. In contrast, NT and AdGFP CMs showed no statistically significant change over the same period. On the other hand, NT CMs treated with 10 µM forskolin increased their beating frequency by 23%. We note that the relative effects of forskolin and light stimulation on beating rates align with our observation of [cAMP]_i_ trends shown in Fig. 2E.

**Fig. 5.**
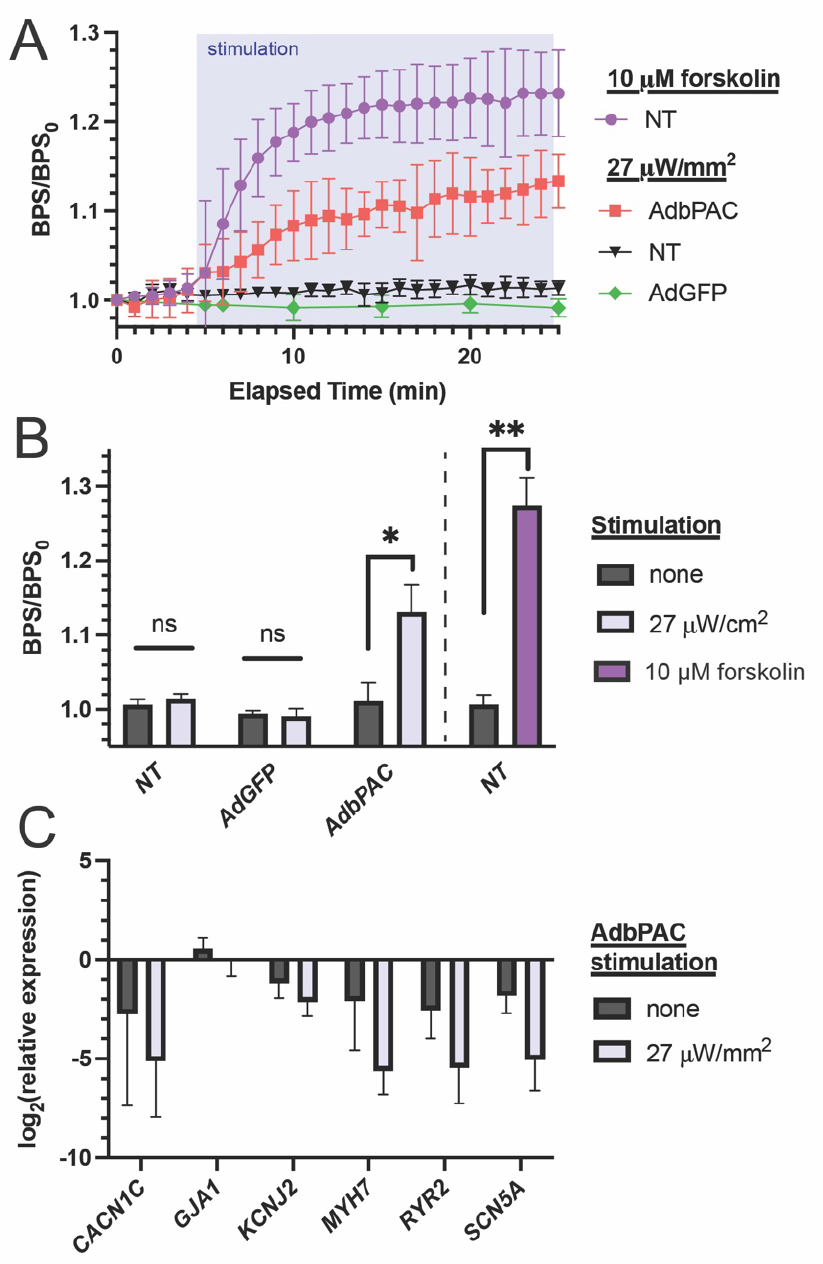
Response to illumination. (A) Comparison of NT, bPAC- and GFP-expressing CMs in response to light (N=4 for each group) or forskolin (N=3). (B) Aggregate endpoint data from panel (A), corresponding to t=25 min (photostimulation for 20 min). Comparisons are by two-tailed t-test, *<0.05, **<0.01. (C) qPCR for key proteins associated with CM contractility. Light stimulation was applied for 30 min. N=3.

The CMs were also examined by quantitative PCR for the expression of various markers relevant to contractibility (*Gja1, Myh7*) and the handling of calcium (*Cacna1c, Ryr2*), potassium (*Kcnj2*) and sodium (*Scn5a*) channels (Fig 5C, Table S1). The level of *Gja1* (encoding for Cx43), was not altered by the expression of bPAC and multi-day, 30-minute illumination, which agrees with the immunofluorescence results (Fig. 4). In addition, no significant suppression of other genes was noted in the bPAC cells kept in dark, suggesting that the functions of CMs remained unchanged with the expression of bPAC. In addition, we observed no statistical difference in the expression of any gene between dark and irradiated groups.

### Irradiation achieves sustained modulation while preserving normal cellular function

We next asked whether photoactivation of bPAC could maintain CMs at a stable, elevated beating rate. To this end, irradiation was applied for 120 minutes. The beating eventually plateaued at a rate 14% greater than the initial rate. Significantly, this increase reached 50% and 90% of maximum by t_50_=7 and t_90_=13 minutes, respectively. By contrast, the basal beating rate of bPAC-expressing CMs in the dark state exhibited no more than 0.04% mean deviation from baseline (Fig. 6A).

**Fig. 6.**
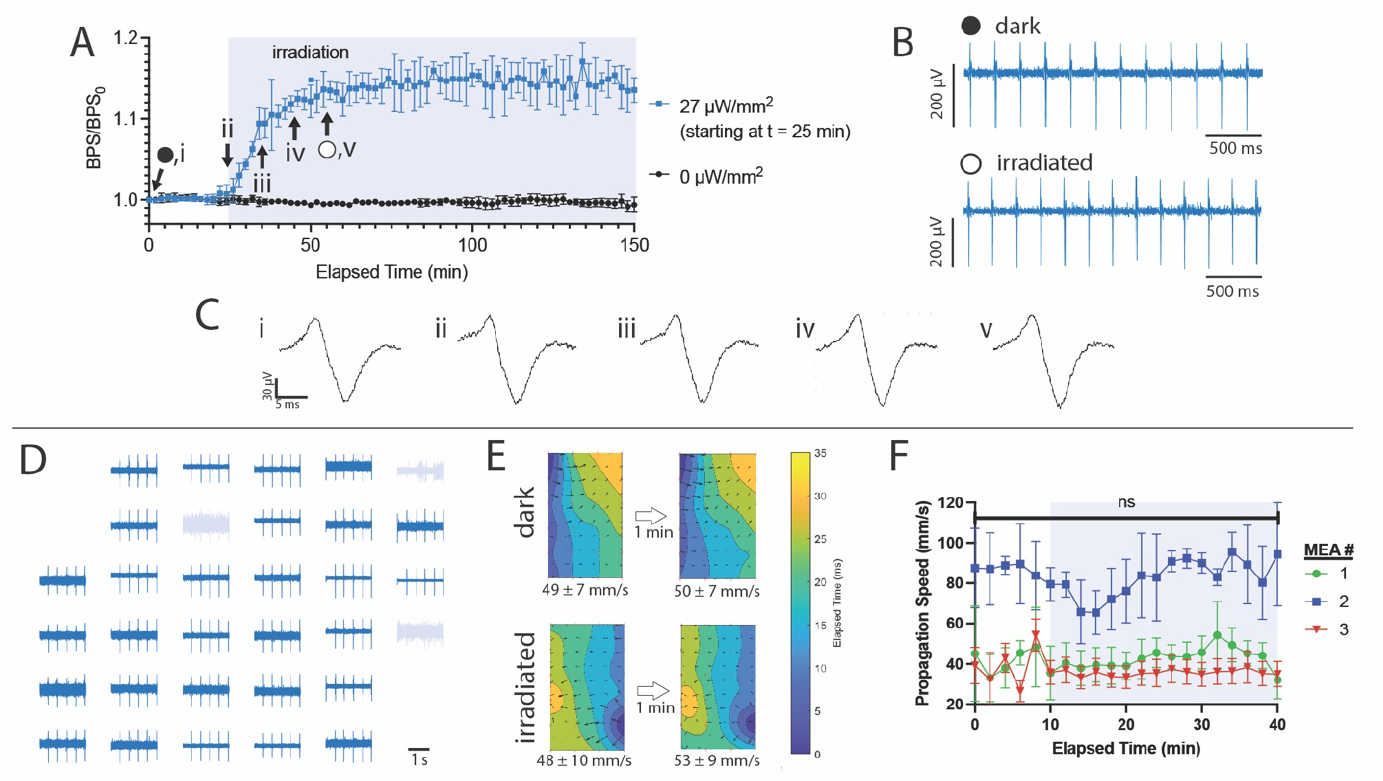
Electrophysiological analysis of optogenetically engineered CMs. (A) Beating rate for chips without (black) or with (blue) 2 hours irradiation starting at t=30 min (light blue region); N=3. (B) Representative single-device readouts in dark and light states corresponding to time points (⠤,◯) noted in panel (A). (C) Representative peak expansions corresponding to time points (i-v) noted in panel (A). (D) Representative readouts from an array of 32 recording elements in the dark state. (E) Isochronal maps representing signal propagation (top) before and (bottom) during illumination. The area represented by each map is 600 µm wide x 1000 µm tall. (F) Propagation speed vs. time during dark and light (light blue region) states on three separate MEA cultures. Error bars represent S.E.M. among all velocity vectors on the isochronal map. Isochronal maps in panel (E) were calculated from data sets for MEA #1.

While the characteristics of extracellular spikes are only weakly correlated to those of the action potential, their magnitude and shape is strongly dependent on the tightness of junctions between the CMs as well as the sheet resistance between the CMs and substrate (*41*). Unaveraged signals from a representative device (Fig. 6B,C) demonstrate uniform shape and magnitude in both dark and light states. These data indicate that irradiation affects neither cell-cell coupling nor the CM adhesion to the substrate. Furthermore, the biphasic shape of these signals can be attributed to stimulus currents injected through gap junctions, followed by inward Na^+^ and Ca^2+^ currents (*42*). The stability of our signals is evidence that these ionic functions are not affected by irradiation.

To quantitatively assess signal conduction through the monolayer we considered multiplexed readouts from the MEA. Figure 6D shows signals from a typical chip with 29 out of 32 functional bioelectronic interfaces. We used a subset of these devices, in this case in a 4x6 rectangular array, to construct isochronal maps, which provide information about wavefront propagation. We obtained readouts throughout a 10-minute dark period followed by 30 minutes of continuous irradiation. Representative maps from dark and irradiated states at 7 DIV showed uniform wavefront propagation patterns across the array. Despite long-term drift, which is expected in the absence of pacing, these patterns were relatively constant over short (1-minute) timeframes (Fig. 6E). Moreover, the average propagation speeds in light (49±7.5 and 50±7.2 mm/s) and dark (48±10 mm/s and 53±9 mm/s) states were not statistically different. These trends were observed across all time points and on three separate chips with CMs from different isolations (Fig. 6F).

The propagation patterns reported here are consistent with those reported for healthy primary CM monolayers (*43*). Uniform signal propagation in space and time is indicative of proper and efficient electromechanical coupling, which is also essential for healthy cardiac function. In contrast, CMs in dysfunctional states presented significantly different propagation characteristics. For example, we previously reported hypoxic CMs with highly nonuniform propagation speeds with averages progressively decreasing over time, highly disordered isochronal maps indicative of multiple activation centers, and patterns changing significantly over ≤1 min time frames (*15*).

### Optical modulation is dose-dependent and time-limited

We hypothesized that we could use the intensity and duration of illumination to customize the modulation of the contractile activity of CMs. To examine the relationship between irradiation intensity and bPAC activity, we irradiated CMs for 30 minutes at intensities ranging from 0 - 27 µW/mm^2^ (Fig. 7A). After 30 minutes, 0, 0.03, 0.3 and 0.7 µW/mm^2^ irradiations yielded progressively larger BPS/BPS_0_ but plateaued at 2.7, 7 and 27 µW/mm^2^ irradiations (p=0.64). These results followed a standard agonist vs. response curve (*44*) with a half maximal effective concentration (EC_50_) of 0.56 µW/mm^2^ (Fig. 7B). They track well with studies in *E. coli*, where [cAMP]_i_ vs. light intensity followed Michaelis-Menten kinetics, with K_m_=3.7±0.4 μW/mm^2^ (*44*).

**Fig. 7.**
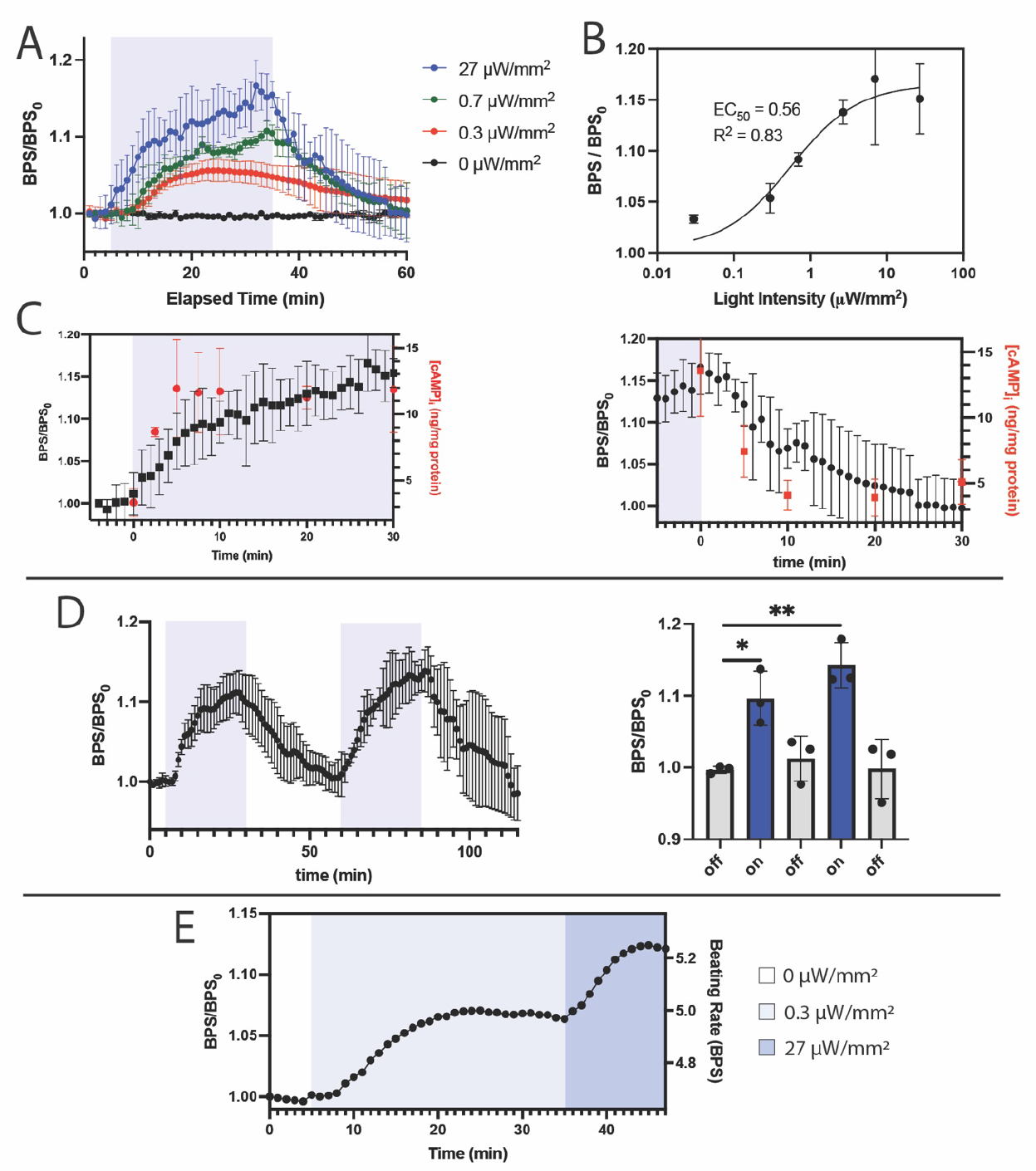
Modulation of contractile activity of CMs with time- and intensity-dependent irradiation. (A) Response to 30-min illumination at selected intensities; N=4 for 27 μW/mm^2^, N=3 for other traces. (B) Summary of beating rates following 30-minute irradiation: regression is agonist vs. response curve. (C) (black) Expansions of rising and falling edges of beating rates after (left) turning the light on and (right) turning the light off. (red) [cAMP]_i_ measured under same conditions. (D) (left) Response to two sequential 30-min illumination cycles; N=3, and (right) corresponding summary statistics; one-way ANOVA; *P<0.05, **P<0.01. (E) Data from a single device in response to stepwise irradiation at 0, 0.3 and 27 μW/mm^2^.

We next considered CM dynamics in response to step changes in irradiation. With 27 µW/mm^2^ intensity, the beating rate reached a plateau at (BPM/BPM_0_)_100_=1.17±0.03. The recorded beating rate closest to 50% of this plateau was (BPM/BPM_0_)_50_=1.08±0.04, at t_50_=6 min (Figure 7C, left). These measurements are in good agreement with the separate experiments shown in Figure 6A, with (BPM/BPM_0_)_100_=1.14±0.02, (BPM/BPM_0_)_50_=1.06±0.1 and t_50_=7 min. Under identical irradiation conditions, [cAMP]_i_ also increased until reaching a plateau, at 13.6±3.5 ng/mg total protein, with t_50_<2.5 min. We observed kinetics with similar trends upon tuning off the light (Figure 7C, right): the elevated beating rate returned to baseline with t_50_=6 min, while [cAMP]_i_ returned to its baseline of 4.2±0.8 ng/mg total protein with t_50_<5 min. These light-modulated CMs exhibited much faster off kinetics than CMs treated with 10 μM forskolin, which after 3 media washes remained at an elevated beating rate with t_50_>5 hours (Fig. S3).

By analogy to remotely-triggered drug delivery systems (*45*), we hypothesized that time- and dose-modulated illumination could be employed to achieve customized beating regimes. We first irradiated CMs over two cycles, with each cycle consisting of a 25-min light phase followed by a 30-min dark phase (Fig. 7D). By the end of the two light phases the CMs reached elevated BPM/BPM_0_ of 1.10±0.04 and 1.14±0.03 (p<0.05 and p<0.01, compared to preceding dark phase). There was no statistical difference between the three dark phases (p=0.79) or the two light phases (p=0.44). In a separate experiment, we irradiated CMs with a stepwise regime consisting of 0, 0.3 and 27 μW/mm^2^ for 5, 30 and 15 minutes, respectively (Fig. 7E). In the representative single-device example shown we found that CMs responded in a stepwise fashion with stable plateaus at 4.67, 5.00 and 5.29 BPS, respectively.

### bPAC expression is stable over multiple days

To assess the stability of the adenovirus and expressed bPAC, we interrogated and irradiated the same MEA on four subsequent days (7-10 DIV), which is an experimental time course representative of other engineered cardiac tissue studies. Upon irradiation, the CM beating rate increased and eventually plateaued on all four days (Fig. 8A). On each of these days, we observed a general, progressive increase in dark-state signal amplitude (Fig. 8B; 44.4, 34.5, 156 and 227 μV from a single, representative device) and baseline beating rate (Fig. 8C; 4.1, 4.0, 5.2 and 5.1 BPS). These trends are consistent with those observed in bPAC-negative CM cultures, which over time develop tighter intercellular and cell/substrate junctions and greater contractility (*10, 46*). Significantly, we also found that BPM/BPM_0_ had a general upward trend of 1.15, 1.16, 1.20 and 1.18 (Fig. 8D).

**Fig. 8.**
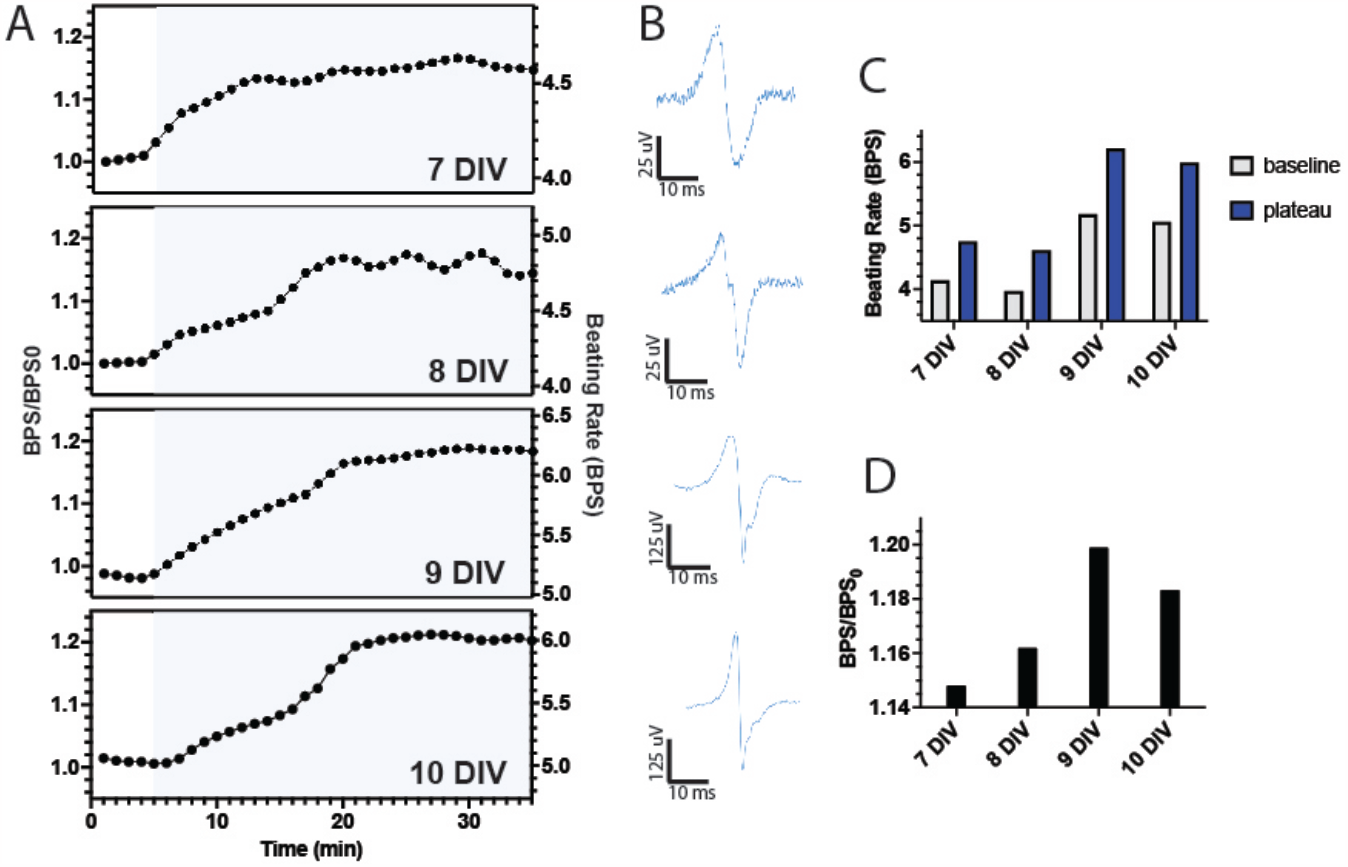
Multi-day studies. (A) BPS/BPS_0_ and beating rate vs. time for a single MEA irradiated on four consecutive days, 7-10 DIV. (B) Baseline signals from a single, representative device on each day. (C) Summary of beating rates at baseline and plateau regions. (D) Summary of BPS/BPS_0_ corresponding to beating rate data shown in panel (C).

## Discussion

We demonstrated an integrated system permitting the modulation of the beating rate of CMs via a photostimulation with concurrent monitoring of their response via multiplexed, bioelectronic readouts. CMs became amenable to altering their contractile activity with light by expressing bPAC, which upon irradiation with blue light increased their [cAMP]_i_ and subsequently the beating rate. The bioelectronic readouts allowed us to continuously measure the beating rate and thus the time-dependent effects of photostimulation. We found that CMs could be maintained at a stable, elevated beating rate for at least two hours. They responded quickly to step changes in irradiation, reaching a new steady state within about 20 minutes. In addition, we found that we could achieve a desired beating rate by adjusting the illumination intensity.

While optical and electronic stimulation approaches have gained wide adoption *in vitro* and *in vivo*, their utility in the clinic has been limited because of the variability in responses across different cell types, between patients, and over time due to tissue plasticity. Bioelectronic therapies that incorporate both stimulation and recording functionalities could address these shortcomings. We demonstrated a customized beating regime by modulating the timing and intensity of photostimulation (Fig. 7E). While that system was open-loop, relying on calibrated beating responses for our specific cell type, we note that both LED light sources and bioelectronic recording elements would be compatible with closed-loop feedback architectures. Such systems have been proposed for neuromodulation (*47*), and could readily be extended to cardiac and other systems. Our optical and bioelectronic interfaces are especially timely given recent advances in machine learning techniques, which are uniquely capable of interpreting large volumes of data to provide real-time interventions, such as the detection and reversal of myocardial ischemia (*48*).

Our modulation of bPAC activity allowed proper control of the CM contractile activity via the cAMP secondary messenger achieving the desired physiological effect. Optogenetic approaches have been developed to regulate gene expression and a wide variety of protein functions, with applications relating to cytokine expression and hemoglobin and insulin production (*49*). Future optical and bioelectronic interfaces could target multiple endogenous pathways and also integrate electrical, chemical and mechanical sensors to achieve regulation in a broad range cell types and biological functions.

Clinical translation of any optogenetic or bioelectronic approach will require that the therapy be administered in a minimally invasive fashion. While the system reported here used a macroscale LED array and rigid bioelectronic device substrate, both technologies have also been demonstrated as microscale arrays on flexible and conformal substrates. Such systems could be implanted onto the surface of an organ, for example the epicardium after surgery (*50*). Very recently, mesh-like bioelectronics were injected into the vascular system and implanted within the brain using an intracortical catheter, precluding invasive surgery altogether (*51, 52*). Additionally, PACs activated by red light (e.g., near infrared window light (*53*)), which exhibits deeper tissue penetration than blue light, could be employed with an extracorporeal light source, thereby eliminating the need for implanting microLED and reducing the implant size. Future versions of our optogenetic and bioelectronic platform could leverage these advances in materials science to achieve stable interfaces with the heart and other regions of the body to address a wide variety of medical conditions.

## Materials and Methods

### Chemicals & Reagents

All reagents used for cardiomyocyte isolation/culture and Pt black deposition were purchased from Sigma-Aldrich (St. Louis, MO): Horse serum, fetal bovine serum, penicillin-streptomycin, aminocaproic acid, ascorbic acid, insulin-transferrin-selenium solution, collagenase II, forskolin, chloroplatinic acid, lead acetate. Four-inch silicon wafers were purchased from University Wafer, and a two-part Sylgard 184 polydimethylsiloxane (PDMS) kit was purchased from Dow Corning (Midland, MI).

### Cardiomyocyte Isolation

We isolated cardiomyocytes from P1 neonatal Sprague-Daley rat pups (Charles River, Wilmington, MA) as described (*54*). Briefly, hearts were isolated and placed in ice-cold PBS glucose solution to remove residual blood and connective tissue. Heart tissue was then minced into ∼1 mm^3^ pieces and suspended in 10 ml of PBS-glucose solution. 7 ml of type II collagenase, warmed to 37°C, was added to the tissue and placed on a tube rotator at 37°C and 5% CO_2_. The tissue suspension was agitated to digest the collagen and release cardiomyocytes into the supernatant. After 7 minutes the tissue was gently titrated 5-6 times and allowed to settle in the conical tube for 3 minutes. The supernatant was then collected and added to 10 ml of DMEM and 10% FBS to inactivate the collagenase. Seven ml of fresh collagenase II solution was added to undigested tissue and this process was repeated until all the collagenase II had been used (7 steps in total). After the final digestion step, the cell solution was filtered through a 70 µm sieve into a new conical tube. The cells were then spun down at 100g for 5 minutes and resuspended in 30 mL of DMEM. To enrich the cardiomyocyte population, the resuspended cells were pre-plated on tissue culture T-175 flasks for 30-45 minutes. Unattached cells were removed and strained again with a 70 µm sieve and spun down at 100g for 5 minutes before cell counting and seeding.

### Cardiomyocyte Cell Culture

Prior to seeding, MEA chips were sprayed down with ethanol and irradiated overnight with UV light. To facilitate cell adhesion, chips were incubated at 37°C with a solution of 1 mg/ml fibronectin dissolved in 0.02% gelatin solution for 1 hour, after which they were ready for seeding. Myocardial culture media was prepared by adding 10% horse serum, 2% fetal bovine serum, 1% penicillin-streptomycin, 1% ITS solution, 6 mg/mL aminocaproic acid, and 50 ug/ml ascorbic acid into DMEM. Cardiomyocytes were seeded onto chips at 2.5x10^5^ cells/cm^2^, maintained in myocardial culture media, and incubated under standard conditions (37°C, 5% CO_2_, 95% humidity) for 10 DIV. Unless stated otherwise, culture media was exchanged daily.

### Generation of adenovirus carrying the bPAC gene and transduction

A human-codon optimized bPAC gene was synthesized (GeneArt, ThermoFisher, Waltham, MA) from the available bPAC sequence (accession number: GU461306.2) and followed with a cMyc-derived epitope, an internal ribosomal entry site (IRES) sequence and mCherry. The sequence of bPAC-cMyc-IRES-mCherry was carried in pShuttle-CMV vector for generation of adenovirus (AdbPAC) in 293AD cells with the AdEasy system (Agilent Technologies, Santa Clara, CA) (*34*). The adenovirus carrying the GFP gene under the CMV promoter (AdGFP) was obtained from the Baylor College of Medicine Vector Development Core (Houston, TX). Purification and titer determination of the adenoviral particles were carried out using the Adeno-X™ Maxi Purification Kit (Takara Bio USA, San Jose, CA) and QuickTiter™ Adenovirus Titer ELISA Kit (Cell Biolabs, San Diego, CA), respectively. The adenovirus propagation was conducted by transducing 293AD cells at 50% confluence with harvested when the cytopathic effect is complete. The cardiomyocytes were transduced at MOI as stated at 24 hours after seeding. Following an incubation period of 48 h with viral particles, fresh medium was replenished, and assays were carried out 24 h after.

### Western blot analysis

Cells were lysed in cell lysis buffer (Cell Signaling Technology, cat. no. 9803, Danvers, MA) supplemented with a protease inhibitor cocktail (Thermo Scientific, cat. no. 78425, Waltham, MA) and total protein was determined (ThermoFisher, cat. no. 23236, Waltham, MA). Cell lysates were boiled for 5 min at 95 °C, loaded to a polyacrylamide gel (30 μg of total protein/lane) and after gel electrophoresis and protein transfer to polyvinylidene difluoride (PVDF) membranes (Millipore, cat. no. IPVH00010, Burlington, MA). The membranes were blocked with 5% milk in Tris-buffered saline with 0.1% Tween-20 (TBST) for 1 h at room temperature. The membranes were incubated with primary antibodies against the c-Myc epitope (Cell Signaling Technology, cat. no. 2278s, Danvers, MA) at 4 °C overnight, or with GAPDH (Sigma Aldrich, cat. no. G9545, St. Louis, MO) for 1 h at room temperature. Following 3 more TBST washes, secondary horseradish peroxidase (HRP)-conjugated antibodies (Jackson ImmunoResearch Laboratories Inc., West Grove, PA) were added for 1 h at room temperature. Membranes were washed 3 times, developed with Clarity Max Western ECL Substrate (Biorad, cat. no. 1705062, Hercules, CA) and detected in a C-DiGit blot scanner.

### Intracellular cAMP measurements

Intracellular cAMP ([cAMP]_i_) was determined 72 h after transduction. One million cells/well were seeded in 12-well plates and 5 wells were used for each condition. One hour before the experiment, 1 mL of fresh medium was added to the wells. Forskolin at 10 µM (PeproTech Inc, Cranbury, NJ) was introduced at the start of the experiment. Cells from non-treated, AdbPAC and AdGFP were incubated with or without illumination for various times at 37 °C and 5% CO_2_. Then, cells were lysed in 0.1 M HCl for determination of [cAMP]_i_ via enzyme-linked immunosorbent assay (ELISA; Cayman Chemical Co., Ann Arbor, MI) following the manufacturer’s instructions. The [cAMP]_i_ concentration was normalized by dividing with the total protein content measured by the Bradford method (ThermoFisher, Waltham, MA).

### Flow cytometry

Harvested cardiomyocytes were treated with TrypLE™ Express Enzyme (1X) (Gibco™, Waltham, MA) for 5 minutes and were centrifuged at 500xg for 5 minutes. The pellet was then resuspended in PBS and after a wash with PBS, cells were incubated with green fluorescent calcein-AM dye (Invitrogen, Waltham, MA) for 20 minutes at room temperature. After the incubation and three washes with PBS, the stained cells were processed by an Attune NxT flow cytometer (Thermo Fisher Scientific, Waltham, MA) to measure calcein-AM and mCherry expression. The results were analyzed with the FCS Express software (v. 6, De Novo Software, Pasadena, CA).

### Immunofluorescence

Cardiomyocytes were seeded on 35 mm glass-bottom culture dishes (Matsunami Glass, Bellingham, WA) coated with fibronectin and gelatin, and were cultured and transduced as described above. The cells were fixed in 4% paraformaldehyde (Millipore-Sigma, Burlington, MA) in PBS for 20 min and permeabilized with saponin (Millipore-Sigma) in PBS for 1 h at room temperature in preparation for staining of nuclear antigens. Samples were washed three times (5 min each time) with PBS under light rotation and blocked with 5% BSA in PBS for 1 h. Antibodies against Connexin-43 (rabbit; Abcam, cat. no. ab11370, Cambridge, MA), α-Actinin (mouse, Sigma, cat. no. A7811, St. Louis, MO), or mCherry (goat, OriGene, cat. no. TA150126, Rockville, MD) were added overnight at 4 °C in 1% BSA in PBS. After three washes with PBS, cells were then incubated with a corresponding secondary antibody conjugated at room temperature for 1 h (anti-rabbit Alexa Fluor 488, anti-mouse Alexa Fluor 647, anti-goat Cy™3, Jackson ImmunoResearch Inc., West Grove, PA) in 1% BSA in PBS. After 3 washes with PBS, nuclear DNA was stained with DAPI (Millipore-Sigma, Waltham, MA) for 10 minutes. Following another three washes in PBS, VECTASHIELD® Antifade Mounting Medium (Vectorlab, Newark, CA) was added to the coverslip. Samples were visualized with a Leica TCS SPE confocal microscope (Leica Microsystems, Wetzlar, Germany).

### RNA extraction, RT-PCR, and quantitative PCR analysis

Cardiomyocytes were isolated from three separate litters and plated on 6-well plates for each condition. Total RNA was extracted using TRIzol (ThermoFisher, Waltham, MA, USA) according to manufacturer’s instructions. Reverse transcription was performed at 70 °C for 5 min and 42 °C for 60 min with 1 µg total RNA using ImProm-II reverse transcriptase (Promega, Madison, WI, USA) and 250 ng oligo(dT)_12-18_ primers (ThermoFisher, Waltham, MA, USA). The resulting complementary DNA (cDNA) and primers were mixed with PowerUp™ SYBR™ Green Master Mix (Applied Biosystems, Waltham, MA), and was analyzed on a StepOne Plus qPCR thermocycler (Applied Biosystems, Foster City, CA, USA) by quantitative PCR (qPCR) for 40 cycles at 58–60°C annealing temperature depending on primer set. Primer sequences are listed in Table S1. Gene expression was analyzed with the ΔΔCT method (*55*). GAPDH served as the endogenous control of gene expression levels.

### MEA Fabrication

MEAs consisted of circular 30-μm diameter gold recording elements and SU-8 passivated interconnects fabricated on a silicon/silicon oxide substrate (*39, 56*). Briefly, photomasks were designed using AutoCAD (Autodesk Inc., San Rafael, CA). The device and interconnect layers were fabricated with LOR 3A and S1813 and metallized with 7 nm Cr / 50 nm Au by sputter deposition (NSC-3000, Nano-Master, Inc., Austin, TX). Interconnects were passivated with 2 µm thick SU-8 photoresist (SU-8 2002, Kayakli Advanced Materials, Westborough, MA). MEA chips were mounted onto a custom printed circuit board (PCB) with 10-pin surface mounted connectors (2.54 mm pitch). Electrical connections between the MEA and PCB were defined with silver epoxy (CW2400, Chemtronics, Kennesaw, GA). A 0.5-in diameter silicone well (VWR, Radnor, PA) was adhered to the chips with PDMS which was cured at 65°C for 4 h. The recording elements were electroplated with platinum black using aqueous chloroplatinic acid (1% w/v) and lead acetate (0.01% w/v) at -0.5 V for 15 s or total charge transfer of q=-100 nC using a Gamry 600+ electrochemical workstation (three electrode setup, Ag/AgCl reference electrode, Pt counter electrode).

### Impedance Measurements

Impedance spectra of recording elements were collected using an electrochemical workstation (Gamry Reference 600+, Gamry Instruments, Inc., Warminster, PA) with a three-electrode setup (Ag/AgCl reference, Pt wire counter) and PBS solvent. We scanned between 10 and 10^6^ Hz with a perturbative potential of 20 mV.

### MEA Measurements

We performed measurements between 6 and 10 DIV. Differential headstage amplifiers (891221, Harvard Apparatus, Holliston, MA) and adapters (890564, Harvard Apparatus, Holliston, MA) were mounted onto the pin connectors and data were collected using a 32-channel recording system (USB-ME32-FAI, Multichannel Systems GmbH, Germany). Signals were amplified 1000x at the headstage, sampled at 25 kSa/s, then digitally filtered post-hoc in MATLAB using a 2^nd^ order Type 1 Chebyshev filter and a 150 -2500 Hz bandpass filter.

### Optical Stimulation

Cells were illuminated with 480 nm light from an LED panel light system (884667106091218, Resurs2 Corporation) mounted parallel to the of the chip at a fixed distance of 5.5 inches. We used an irradiation measurement probe (PM100D, Thorlabs Inc., Newton, NJ) to measure irradiance. For intensity variation experiments we attenuated the light with combinations of 1.0, 0.6, and 0.3 neutral density filters (53-705, 53-704, 53-703, Edmund Optics, Barrington, NJ).

### Beating Rate and Isochronal Maps

Segments of filtered data (6s length) were processed with custom scripts in MATLAB (MathWorks Inc., Natick, MA) as described (*56*). Beat rates and amplitudes were calculated by peak identification. BPS_0_ represents the beat rate of the first segment in a time series and was used to normalize all subsequent recordings in the series.

Isochronal maps were generated by identifying relative time shifts among all functioning devices on the array. These activation times were interpolated and their gradient (∇ = **i**∂/∂x + **j**∂/∂y) was taken to generate a matrix of vectors representing the local slope of the isochronal map. Local propagation speeds were found by taking the inverse of each vector. Reported propagation speeds represent the average over all vectors on a single chip.

### Statistical Analysis

All statistical analyses were performed in GraphPad Prism (GraphPad Software, San Diego, CA). Groups were compared using student’s t-test or two-way ANOVA, and are presented as mean ± standard deviation, unless otherwise specified. A p< 0.05 was considered statistically significant.

## Acknowledgments

We thank C. Fucetola and J. Wang for helpful discussions. This work was performed in part at the Tufts Micro and Nano Fabrication Facility. We thank J. Vlahakis for assistance with fabrication processes. This work was also performed in part at the Harvard

University Center for Nanoscale Systems (CNS); a member of the National Nanotechnology Coordinated Infrastructure Network (NNCI), which is supported by the National Science Foundation under NSF award no. ECCS-2025158.

## Author contributions

Conceptualization: OAB, ZC, EST and BPT.

Preparation of adenoviruses, gene transduction, Western blot and ELISA: ZC Immunofluorescence: ZC

Cardiomyocyte isolation: OAB and Y-RL

Bioelectronics measurements: OAB

Device fabrication and assembly: OAB, MC, AAR, HL

Data analysis: OAB, ZC, EST and BPT

Writing—original draft: OAB, ZC, EST and BPT

Writing—review & editing: all authors

## Data and materials availability

All data are available in the main text or the supplementary materials.

