## Supplementary Information for "An Integrated Optogenetic and Bioelectronic Platform for Regulating Cardiomyocyte Function"

Olurotimi A. Bolonduro *et al.*

**This PDF file includes:**

Figs. S1 to S3
Table S1

**Fig. S1.**

Assembly consisting of PCB, MEA chip and silicon well.

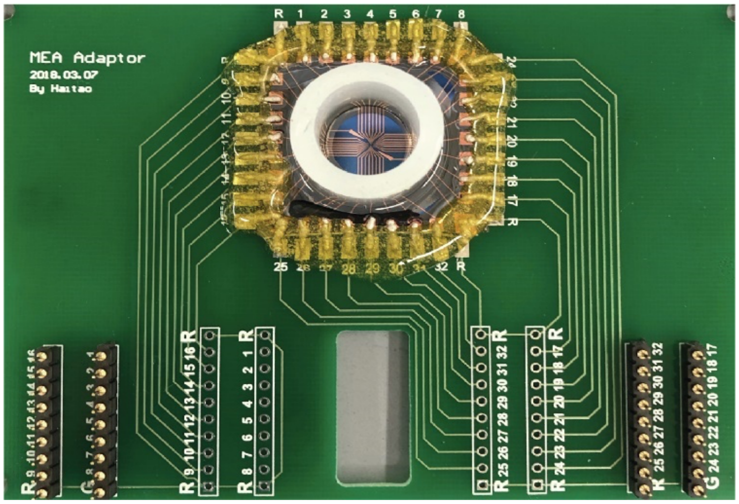

**Fig. S2.**

(a,b) Impedance and phase spectra of (magenta) bare gold and (black) Pt black-coated electrodes. (c) Impedance at 1 kHz. N=20, **** p<0.0001.

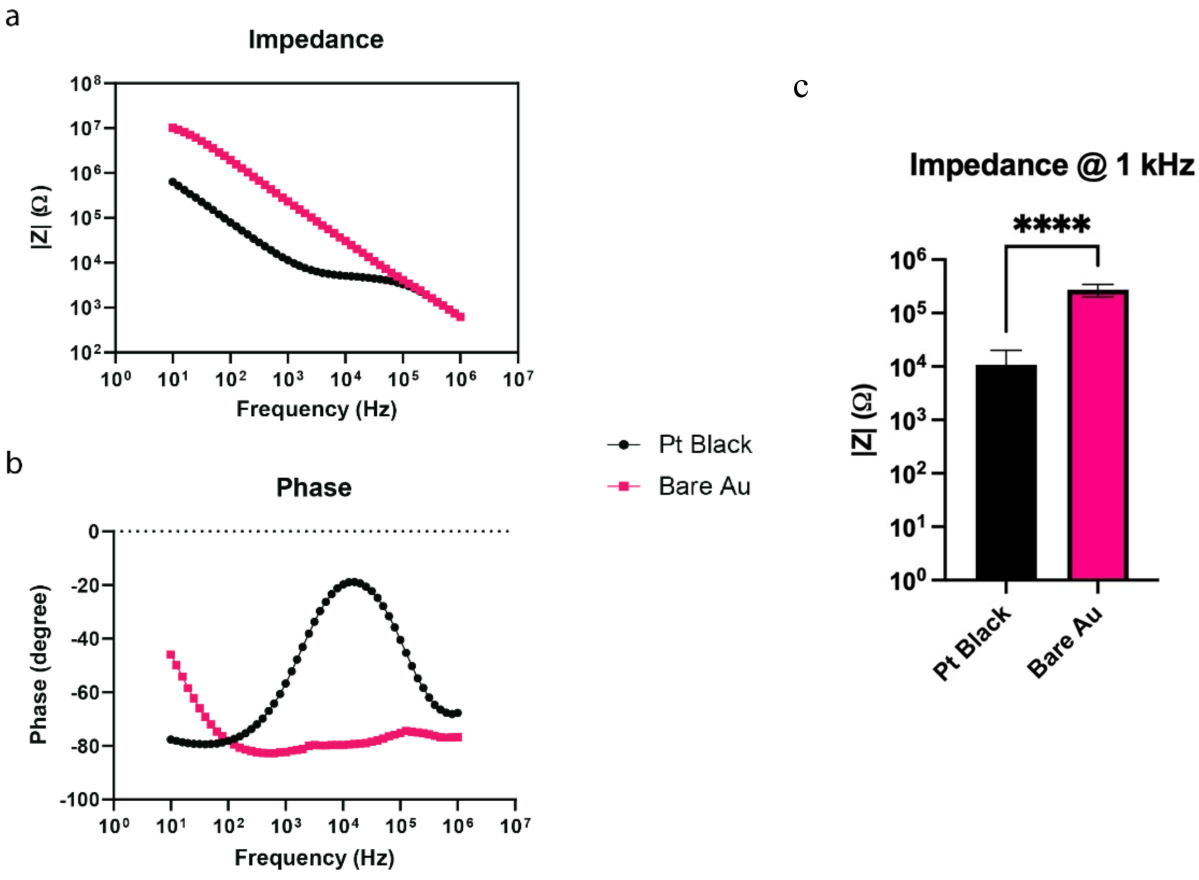

**Fig. S3.**
Normalized beating rate after stimulation with (black) light and (magenta) forskolin.

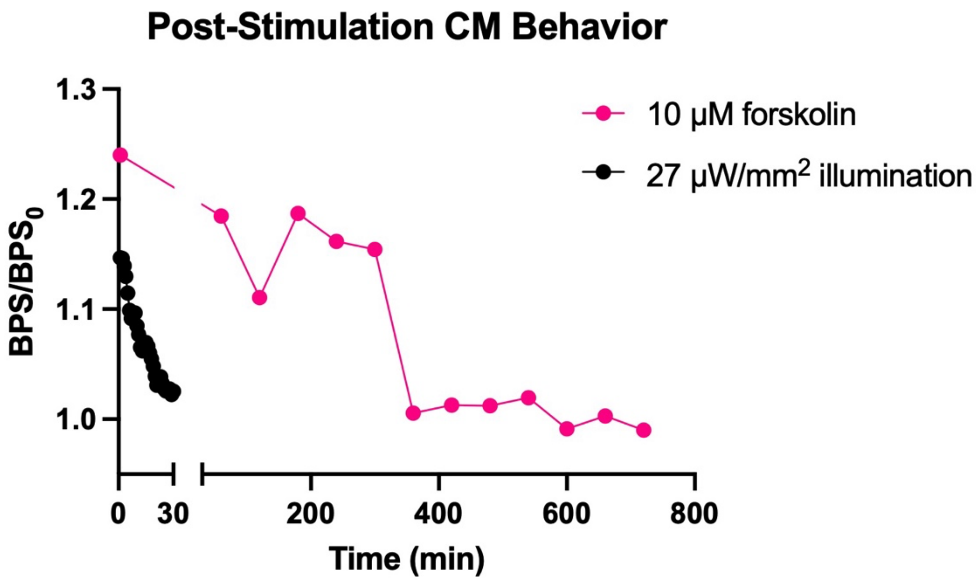

**Table S1.**
Primers used for qPCR assays in this study (shown in a 5′-to-3′ orientation)

| Gene | Amplicon Size (bp) | Primers | |
|---|---|---|---|
| ***Ryr2*** | 99 | Forward | ACTGCTGGGCTACGGCTAC |
| | | Reverse | CTGAAGATGCGGAACCTCTC |
| ***Cacna1c*** | 109 | Forward | GTTGCCCTGGGTGTATTTTG |
| | | Reverse | GGCTTTCTCCCTCTCTTTGG |
| ***Kcnj2*** | 116 | Forward | GCACAAGTACGGACTCACCT |
| | | Reverse | TCCAAAGACAGAATCGGCCA |
| ***Gja1*** | 99 | Forward | CGCCGGCTTCACTTTCATTA |
| | | Reverse | GGTGGAGTAGGCTTGGACCT |
| ***Myh7*** | 145 | Forward | TGGCACCGTGGACTACAATA |
| | | Reverse | TACAGGTGCATCAGCTCCAG |
| ***Gapdh*** | 92 | Forward | GACATGCCGCCTGGAGAAAC |
| | | Reverse | AGCCCAGGATGCCCTTTAGT |
| ***Scn5a*** | 220 | Forward | ACATGTTCAACTTCCAGACCTTC |
| | | Reverse | ACGATGAGGAAGGAGATGATGAT |
